# Accurate and Reproducible Whole-Genome Genotyping for Bacterial Genomic Surveillance with Nanopore Sequencing Data

**DOI:** 10.1101/2025.02.24.639834

**Authors:** K. Prior, K. Becker, C. Brandt, A. Cabal Rosel, J. Dabernig-Heinz, C. Kohler, M. Lohde, W. Ruppitsch, F. Schuler, G.E. Wagner, A. Mellmann

## Abstract

Despite recent advances in error rate reduction, until recently Oxford Nanopore Technology (ONT) sequences lacked the accuracy required for fine scale bacterial genomic analysis. Here, recent software improvements of ONT and the ONT-cgMLST-Polisher within the SeqSphere^+^ software were evaluated.

We used short-(Illumina) and long-read ONT sequences of 80 multidrug-resistant bacteria (MDROs) for benchmarking. Illumina reads were *de-novo*-assembled using SKESA. For ONT, Dorado super accurate (SUP) model 4.3 or 5.0 basecalled reads were assembled with Flye and then polished with Medaka v1.12 m4.3 or Medaka v2.0 bacterial methylation model. In addition, the ONT-cgMLST-Polisher was run over all assemblies. The ‘ground truth’ (GT) hybrid assemblies were created using Hybracter v0.10.0. Sixteen isolates from four species out of the original 80 isolates were sent to six laboratories for a ring trial.

The 80 MDROs basecalled with SUP m4.3 had an average cgMLST allele distance (AD) to the GT of 4.94 with Medaka v1.12 and 1.78 with Medaka v2.0, respectively. After further polishing the Medaka v2.0 data with the ONT-cgMLST-Polisher, the AD dropped to 0.09. Using data basecalled with SUP m5.0 with Medaka v2.0 further reduced the AD significantly to 0.04. While the ring trial data basecalled with Dorado SUP m4.3 showed more variability and insufficient results for some samples, model 5.0 data resulted in average ADs of 0.36 and 0.17 without and with the ONT-cgMLST-Polisher, respectively. In conclusion, recent ONT Dorado and Medaka models combined with the ONT-cgMLST-Polisher improved ONT sequencing accuracy and made it sufficiently reproducible for genomic surveillance of bacteria.

**Importance**

Oxford Nanopore Technologies (ONT) sequencing methodology is especially attractive for small and medium-sized laboratories due to its relatively low capital investment and price per sample consumable costs. However, until recently it lacked accuracy and reproducibility for bacterial genomic genotyping. Here, we present an evaluation of the most recent ONT bioinformatic (basecalling and polishing of consensus) improvements and a new ONT-cgMLST-Polisher tool. We demonstrate that by applying those procedures ONT whole-genome genotyping-based surveillance of bacteria is finally accurate and reproducible enough for routine application even in small laboratories

## Introduction

Whole-genome sequencing (WGS)-based genotyping is an essential part of infectious disease control (1). However, reliable results require a low error-rate and high reproducibility. Short-read sequencing (i. e., Illumina technology [Illumina Inc., San Diego, CA, USA]) has been the gold standard for many years due to its high accuracy, but lacks the ability to fully reconstruct mobile genetic elements, which play a major role in the dissemination of antimicrobial resistance (AMR) (2, 3). Long-read sequencing overcomes this limitation, however, until recently, the available technologies were either hampered by the capital investment for routine laboratories (PacBio sequencing [Pacific Biosciences Inc., Menlo Park, CA, USA]) or did not provide the required accuracy (Oxford Nanopore sequencing [Oxford Nanopore Technologies, Oxford, UK]). However, recent advances in Oxford Nanopore Technologies (ONT) sequencing chemistry and flow cells as well as basecalling models significantly reduced their error rates. In 2022, Sereika *et al.* (4) reported that the quality of assemblies of ONT R10.4 sequences was comparable to Illumina data. Similarly, a study comparing Illumina and ONT sequencing of 356 multidrug-resistant organisms (MDRO) found ONT sequence accuracy generally suitable for genomic surveillance with the exception of *Pseudomonas aeruginosa* (5). In contrast, Lerminiaux *et al.* (6) found ONT R10.4 assemblies to be only sufficient for plasmid characterization and AMR detection but still recommended short-read polishing for fine-scale analyses such as single nucleotide polymorphisms (SNP) detection. Another study found extensive base errors caused by methylation sites, which led to discrepancies in core-genome multi-locus sequence typing (cgMLST) and resulted in false exclusions from outbreak clusters in *Klebsiella pneumoniae* (7).

In a multicenter study, Dabernig-Heinz *et al.* (8) analyzed cgMLST results of four public health-relevant species and found highly strain-specific typing errors related to consistent DNA methylation motifs. Again, these errors led to discrepancies in cgMLST outbreak clusters in some cases. The authors were able to improve the results by PCR preamplification, more recent basecalling models and optimizing the polishing step. However, they concluded that improvements are still necessary to gain the accuracy needed for bacterial genomic surveillance and that computational approaches have the potential to deliver these improvements if strain-specific differences are taken into account when training new models.

Since this study was conducted, several advances in the bioinformatic tools used for processing ONT data became available. First, a new basecalling model, Dorado SUP m5.0, was published, which increases the accuracy compared to Dorado SUP m4.3 but is much slower and not (yet) part of MinKNOW, the software which controls the sequencing and enables real-time basecalling. Further, with Medaka v2.0, a new polishing model was introduced, which was specifically trained to reduce errors resulting from bacterial methylation. Finally, SeqSphere^+^, a widely used software for cgMLST-based bacterial genotyping (9), includes a new ONT-cgMLST-Polisher.

To determine the remaining error rates, we sequenced 80 MDRO including all ESKAPE species (*Enterococcus faecium, Staphylococcus aureus, Klebsiella pneumoniae, Acinetobacter baumannii, Pseudomonas aeruginosa, and Enterobacter species*), the leading causes of nosocomial infections throughout the world, with Illumina and ONT and evaluated the various bioinformatics solutions mentioned above. Further, to analyze the reproducibility, we conducted a ring trial with six laboratories that were sent DNA from 16 challenging samples of this set of microorganisms.

## Materials and Methods

### Strains and DNA isolation of evaluation dataset

A total of 80 MDRO belonging to eleven species (Table S1) were grown overnight on Columbia Agar at 35±2°C (ambient air). Genomic DNA (gDNA) was extracted using the ZymoBiomics DNA Miniprep Kit (Zymo Research, USA) following the manufacturer’s instructions. Quantity and quality of the extracted gDNA were assessed using the Qubit 2.0 dsDNA BR Kit (Thermo Fisher Scientific, USA) and a NanoDrop spectrophotometer (Thermo Fisher Scientific, USA), respectively.

### Short read sequencing

Illumina Library was prepared with the Illumina DNA Prep Kit and UDI Indexes according to the manufacturer’s instructions (Illumina Inc.). Sequencing was performed using an Illumina NextSeq 2000 platform (Illumina Inc.) generating 2×100 bp paired-end reads. Adapter-trimmed FASTQ files were downsampled to 100x and *de novo* assembled with SKESA v2.3.0 (10) in combination with BWA-MEM v0.7.15 (11) for read mapping and consensus calling with no coverage minimum. A consensus call read support threshold of 60% was applied.

### Long-read sequencing

The ONT MinION in combination with the v14 chemistry (SQK-RBK114.24) was used for long-read sequencing. Library preparation was conducted with the Rapid Barcoding Kit v14 (RBK, ONT) and basecalling was performed using Dorado v0.7.1 with super accurate mode (SUP) models version 4.3 (ONT, dna_r10.4.1_e8.2_400bps_sup@v4.3.0) and version 5.0 (ONT; dna_r10.4.1_e8.2_400bps_sup@v5.0.0). Chopper v0.7.0 (12) was used for read trimming with the following parameters: quality threshold of 10 and minimum read length of 500 bp. Reads were downsampled to 100x coverage in deterministic mode using Rasusa v0.8.0 (13). Flye v2.9.3-b1797 (14) was used for *de novo* assembly with the parameters --nano-hq --deterministic. Medaka v1.12 (ONT) in combination with the fitting version 4.3 polishing model and the bacterial methylation model of Medaka v2.0.1 (ONT; https://github.com/nanoporetech/medaka) were applied for assembly polishing.

### Ground truth assemblies

Hybrid assemblies representing the ‘ground truth’ (GT) for all comparisons were generated using Hybracter v0.10.0 (15) and Medaka v2.0.1 with the bacterial methylation model, utilizing ONT SUP model m4.3 datasets combined with the Illumina short-read data.

### cgMLST data analysis

All assemblies were analyzed with SeqSphere^+^ version 10.5 (Ridom GmbH, Münster Germany) using the available public cgMLST schemes (https://www.cgmlst.org/) for the following species: *Acinetobacter baumannii, Enterococcus faecium, Escherichia coli, K. pneumoniae, P. aeruginosa, Serratia marcescens,* and *Staphylococcus aureus.* New public cgMLST schemes were generated for *Enterobacter hormaechei*, *Morganella morganii*, and *Proteus mirabilis*, while a stable but not public scheme with 2,839 cgMLST targets was created for *Citrobacter freundii*.

### Priority AMR target analysis

The NCBI AMR Finder Plus v3.11.26 (16) task template included in the SeqSphere^+^ software was used to find specific genes from the NCBI Bacterial Antimicrobial Resistance Reference Gene Database. Targets that might confer resistance to carbapenems, colistin, vancomycin, or methicillin or that contain ESBL or AmpC in their name were highlighted as priority AMR targets.

### ONT-cgMLST-Polisher

All ONT datasets were analyzed with and without the proprietary ONT-cgMLST-Polisher v1.0 that is part of the SeqSphere^+^ v10.5 ONT Data Assembly module. This tool maps Dorado SUP basecalled FASTQ reads (≥ model 4.2) to a Medaka-derived assembly consensus FASTA sequence using Minimap2 (17). The alignment is then scanned for potential methylation-related sequencing errors in core and accessory genome MLST genes, e. g., differing strand-specific majority consensus calls. Those ‘ambiguous’ positions are then compared against a sequence with a closely related cgMLST allelic profile. Finally, based on the comparison the consensus sequence of ambiguous positions is either confirmed or masked with an ‘N’ call. Thereby, core genome MLST target genes that contain an ‘N’ are regarded as missing targets as they fail the quality control that does not allow for ambiguous bases by default.

### Strain selection and procedure for the ring trial dataset

For the ring trial, three Gram negative and one Gram positive bacterial species of the evaluation dataset (*K. pneumoniae, E. hormaechei, M. morganii, and E. faecium*) with chromosome sizes of approximately 5.7, 4.9,4.1, and 2.9 Mb, respectively, were selected focusing on isolates exhibiting large distances to the GT. From each species, four isolates were chosen, including both high- and low-distance samples (Table S1, isolates in bold). This selection included the five ‘worst’ isolates. To rule out the possibility that the allelic differences were a result of contamination, the datasets of the selected isolates were analyzed with CheckM2 (18). Cultivation, isolation, and quality control of the isolated gDNA was performed as described for the evaluation data set. Aliquots from the same DNA preparations were sent to the six participating laboratories.

Library preparation was performed with RBK v14 chemistry according to the manufacturer’s recommendations. Three laboratories used a fixed 200 ng gDNA input per sample, whereas the other three adjusted input according to genome size: 150 ng for ∼3 Mb genomes, 200 ng for ∼4 Mb genomes, 250 ng for ∼5 Mb genomes, and 300 ng for ∼6 Mb genomes. All participating laboratories performed one ONT MinION or GridION sequencing run for 72 hours using an ONT FLOMIN-114 flow cell (R10.4.1 pores). Initial basecalling was done using the Dorado server included in MinKNOW v24.06 in SUP mode with the default model 4.3 and a minimum Q-score of 10. Re-basecalling of raw data was subsequently performed using Dorado v0.8.3 with the SUP m5.0. Following basecalling, all datasets underwent the same analysis pipeline described for the evaluation dataset with two exceptions. Datasets were not downsampled prior to assembly and only the bacterial methylation model of Medaka v2.0.1 was used for polishing. Each dataset was analyzed with and without the ONT-cgMLST-Polisher and compared against the GT.

### Data Availability

Illumina and ONT read datasets of the evaluation data set as well as of the ring trial data of all participants have been deposited under Study accession nos. PRJEB86764 (evaluation dataset) and PRJEB86767 (ring trial data) in the European Nucleotide Archive (ENA), respectively.

## Results

### Evaluation dataset cgMLST error analysis

In total, 80 isolates were sequenced with Illumina and ONT, with all datasets achieving a minimum sequencing depth of > 50-fold, using a *de novo* hybrid assembly approach thereby generating the GT for all subsequent comparisons. Of the Illumina data, only one sample deviated by a single allele from the GT (Table 1). The ONT data generated with the Dorado SUP m4.3 were subjected to several analysis pipelines. A Flye-only assembly, without any polishing, resulted in 37 samples exhibiting differences to the GT, with a maximum allele distance (AD) of 98 alleles (range: 0-98, median, 0). Additional polishing with Medaka v1.12 reduced the number of deviating samples to 33, although the maximum AD remained at 98. Further improvement was observed after polishing the Flye assembly with Medaka v2.0, where only 13 samples showed a deviation from the GT and the maximum AD was reduced to 57 (range: 0-57, median: 0). Moreover, incorporating the ONT-cgMLST-Polisher into the workflow led to a substantial reduction in ADs - yielding maximum ADs of 4 for both the Flye-only (median: 0) and Flye plus Medaka v1.12 pipelines (median: 0), and 3 for the Flye plus Medaka v2.0 pipeline (median: 0). Notably, when used without the ONT-cgMLST-Polisher, Medaka v2.0 reduced the average cgMLST AD from approximately 4.5 (Flye only) to 1.8, whereas its combination with the ONT-cgMLST-Polisher lowered this average to 0.09.

**TABLE 1.** Error analysis results of the 80 sequenced isolates from the evaluation.

|  |  | Samples with<br>Distance to GT | Max. cgMLST<br>Allele Distance | Avg. cgMLST<br>Allele Distance<br>(SD) | Avg. Missing<br>cgMLST Alleles<br>(SD) | Max. Missing<br>cgMLST Alleles | Avg. Ns Called |
| --- | --- | --- | --- | --- | --- | --- | --- |
|  | GT |  |  |  | 17.14<br>(16.93) | 103 |  |
|  | Illumina | 1 | 1 | 0.01 (0.11) | 19.81<br>(17.29) | 109 |  |
| Dorado SUP m4.3 | Flye only | 37 | 98 | 4.53<br>(14.48) | 19.34<br>(17.42) | 103 | n.a.* |
|  | Medaka v1.12 SUP 4.3 model | 33 | 98 | 4.94 (14.9) | 19.06<br>(17.58) | 103 | n.a. |
|  | Medaka v2.0 bacterial methylation model | 13 | 57 | 1.78 (8.17) | 18.11<br>(16.94) | 103 | n.a. |
|  | Flye only + ONT-cgMLST-Polisher | 6 | 4 | 0.15 (0.62) | 26.00<br>(26.59) | 152 | 18.75 |
|  | Medaka v1.12 SUP 4.3 model + ONT-cgMLST-Polisher | 7 | 4 | 0.16 (0.60) | 26.11<br>(27.15) | 149 | 20.14 |
|  | Medaka v2.0 bacterial methylation model + ONT-cgMLST-Polisher | 5 | 3 | 0.09 (0.40) | 22.13<br>(20.76) | 104 | 12.13 |
| Dorado SUP m5.0 | Flye only | 20 | 10 | 0.76 (2.00) | 17.56<br>(16.87) | 103 | n.a. |
|  | Medaka v1.12 SUP 4.3 model | 23 | 8 | 0.73 (1.60) | 17.55<br>(16.89) | 103 | n.a. |
|  | Medaka v2.0 bacterial methylation model | 3 | 1 | 0.04 (0.19) | 17.30<br>(16.91) | 103 | n.a. |
|  | Flye only + ONT-cgMLST-Polisher | 4 | 1 | 0.05 (0.22) | 19.65<br>(16.53) | 105 | 7.96 |
|  | Medaka v1.12 SUP 4.3 model + ONT-cgMLST-Polisher | 2 | 1 | 0.03 (0.16) | 19.64<br>(16.47) | 105 | 9.25 |
|  | Medaka v2.0 bacterial methylation model + ONT-cgMLST-Polisher | 2 | 1 | 0.03 (0.16) | 18.70<br>(16.56) | 105 | 7.14 |

dataset basecalled with Dorado SUP models 4.3 and 5.0
\* n.a. – not applicable

The ONT-cgMLST-Polisher masks ambiguous positions by substituting them with an ‘N’. In the Dorado SUP m4.3 datasets, this resulted in an average of 18.75 Ns for the Flye-only assemblies, 20.14 Ns for the Flye plus Medaka v1.12 assemblies, and 12.13 Ns for the Flye plus Medaka v2.0 assemblies.

Data obtained using the ONT Dorado SUP m5.0 model (version 0.8.3) - where basecalling takes two to three times longer than with model m4.3 and currently does not support live basecalling in MinKNOW - showed a dramatic reduction in both the number of samples deviating from the GT and the maximum cgMLST AD across all analysis setups. When combined with the ONT-cgMLST-Polisher, the maximum AD was reduced to one in every setting. The average cgMLST AD was lowered to 0.05 for the Flye-only assembly and to 0.03 when Medaka v1.12 or Medaka v2.0 was additionally employed. Moreover, the average number of Ns introduced by the ONT-cgMLST-Polisher was lowest (7.14) in the Flye plus Medaka v2.0 pipeline and highest (9.25) in the Flye plus Medaka v1.12 pipeline but in all m5.0 setups approximately 50% lower than the values observed in the m4.3 datasets.

### Evaluation dataset priority AMR target analysis

In our study, the recovery and localization of priority AMR targets was evaluated by comparing Illumina short-read data with ONT data processed using the SUP 4.3 and SUP 5.0 basecalling models against the GT (Table 2). The ONT datasets demonstrated excellent performance with nearly 99% of AMR targets correctly placed either as chromosomal or plasmid-borne, while Illumina data achieved a correct placement rate of only 84%. When assessing misassignments on plasmids, ONT SUP m4.3 showed the best performance with only 3% of targets wrongly placed, followed by ONT SUP m5.0 at 6% and Illumina at 9%. The misplaced targets were mainly genes encoding for resistance against beta-lactams or vancomycin. Furthermore, ONT data did not exhibit any missing AMR targets, whereas Illumina results lacked 6% of the expected targets. The number of excess AMR targets was also lowest with ONT SUP m4.3 compared to both ONT SUP m5.0 and Illumina. Finally, while only 6% of the samples processed with both ONT basecalling model showed errors, Illumina short-read sequencing was erroneous in 19% of the samples.

**TABLE 2.** Priority AMR target recovery and chromosomal or plasmid-borne location results of the evaluation dataset (n=80 multidrug-resistant isolates). Total numbers as well as the numbers per specific target or species are given. Polishing of ONT data was performed with Medaka v2 bacterial methylation model.

|  | Illumina | ONT SUP m4.3 | ONT SUP m5.0 |
| --- | --- | --- | --- |
| **Correctly placed AMR targets** | **136 (84%)** | **160 (99%)** | **161 (99%)** |
| **AMR targets incorrectly placed on chromosome** | **2 (1%)**<br>- blaCTX-M-1 (n = 1)<br>- blaVIM-1 (n = 1) | **0 (0%)** | **0 (0%)** |
| **AMR targets incorrectly placed on plasmid** | **15 (9%)**<br>- blaCMY-2 (n = 1)<br>- blaCTX-M-15 (n = 2)<br>- blaCTX-M-2 (n = 1)<br>- blaOXA-23 (n = 2)<br>- blaSHV-2 (n = 1)<br>- blaVEB-6 (n = 1)<br>- vanB cassette (n = 7) | **5 (3%)**<br>- blaCTX-M-15 (n = 2)<br>- blaSHV-12 (n = 1)<br>- blaSHV-2 (n = 2) | **10 (6%)**<br>- blaCTX-M-15 (n = 1)<br>- blaDHA-1 (n = 1)<br>- blaSHV-2 (n = 1)<br>- vanA-cassette (n = 7) |
| **Missing AMR targets** | **9 (6%)**<br>- blaCTX-M-15 (n = 2)<br>- blaSHV-2 (n = 1)<br>- blaOXA-23 (n = 4)<br>- blaOXA-35 (n = 1)<br>- mcr9.2 (n = 1) | **0 (0%)** | **0 (0%)** |
| **Excess AMR targets** | **0 (0%)** | **3 (2%)**<br>- blaCTX-M-15 (n = 2)<br>- blaSHV-12 (n = 1) | **9 (6%)**<br>- blaCTX-M-15 (n = 1)<br>- blaDHA-1 (n = 1)<br>- vanA-cassette (n = 7) |
|  | Illumina | ONT SUP m4.3 | ONT SUP m5.0 |
| **# of samples with errors (n = 80)** | **15 (19%)**<br>- A. baumannii (n = 2)<br>- C. freundii (n = 1)<br>- E. coli (n = 1)<br>- E. faecium (n = 1)<br>- K. pneumoniae (n = 4)<br>- P. mirabilis (n = 4)<br>- P. aeruginosa (n = 1)<br>- S. marcescens (n = 1) | **5 (6%)**<br>- E. coli (n = 2)<br>- K. pneumoniae (n = 3) | **4 (6%)**<br>- E. faecium (n = 1)<br>- K. pneumoniae (n = 3) |

Priority AMR Targets (in total 162 in GT, i.e., Hybracter 10.0.0 hybrid assemblies with Medaka v2.0.1 from ONT SUP m4.3) are targets that might confer resistance to carbapenem, colistin, vancomycin, methicillin, or that contain ESBL or AmpC in their name. All genomes of the n=80 isolates with Illumina and ONT coverages >50x. Illumina and ONT with SKESA or Flye assembled, respectively.

### Ring trial run statistics

The total number of called bases ranged from 5.83 to 17.02 Gb for the six labs, while the N50 values varied between 6.35 and 9.91 kb, and the average depth of coverage assembled was between 78 and 266 (Table 3). Only in one lab, four out of the 16 samples did not achieve the recommended sequencing depth of 50-fold (https://a.storyblok.com/f/196663/x/f90d1fe1a4/workflow-bacterial-isolates.pdf). Interestingly, neither the number of total (range 978-1.528) nor the number of sequencing (442–511) pores at the start correlated with the output. However, a higher number of total and sequencing pores after 90 minutes did correspond to a higher final output. In the present study, three participating labs applied genome size compensation (GSC) to account for differences in bacterial genome sizes and to normalize coverage distribution across sequencing runs. In general, the average coverage assembled ranged from 78-fold to 266-fold. Notably, except for Lab 1, which exhibited an overall lower coverage, GSC led to a more even coverage distribution, as demonstrated by the narrower standard deviation of the datasets where GSC was applied (Table 3).

**TABLE 3.** Run statistics and coverage metrics of the ring trial data (16 isolates per laboratory).

| Lab no. | Instrument | Genome size comp. | Called bases [Gb] | N50 [kb] | No. of pores at start (total/sequencing) | No. of pores after 1h30min (total/sequencing) | Average Coverage Assembled (SD) |
| --- | --- | --- | --- | --- | --- | --- | --- |
| Lab 1 | MinION | No | 5.83 | 6.35 | 1361/503 | 797/387 | 78 (40.50) |
| Lab 3 | GridION | No | 15.35 | 6.69 | 1528/511 | 1539/505 | 266 (138.85) |
| Lab 4 | MinION | No | 17.02 | 9.91 | 1083/476 | 1224/487 | 233 (106.86) |
| Lab 2 | MinION | Yes | 11.01 | 8.98 | 1453/505 | 1372/498 | 155 (75.18) |
| Lab 5 | MinION | Yes | 14.39 | 8.13 | 978/442 | 1074/459 | 198 (77.81) |
| Lab 6 | GridION | Yes | 16.06 | 8.46 | 1249/488 | 1250/483 | 215 (73.69) |

A total of four samples (Lab 1) did not achieve the desired coverage of >50x: 1x *E. hormachei* (35x), 2x *K. pneumoniae* (44/45x), 1x *M. morganii* (46x)

### Ring trial cgMLST error analysis

Evaluation datasets of the 16 selected ring trial isolates were examined with CheckM2 to assess potential contamination. The analysis revealed that 11 isolates exhibited no contamination (< 0.5% contamination) whereas five samples (A24994, A26728, A27739, A31008, A33124) showed only low-level contamination (< 5%), indicating that discrepancies from the GT could not be attributed to gDNA contamination.

To analyze the reproducibility using the different software tools, we compared the cgMLST ADs of the 16 ring trial samples against the GT for each participating lab. We analyzed the results of data basecalled with Dorado SUP m4.3 (Table 4a) and Dorado SUP m5.0 (Table 4b), as well as with and without the ONT-cgMLST-Polisher. The detailed results, broken down by sample ID and participant, can be found in Table S2. For the data basecalled with Dorado SUP m4.3, the mean average cgMLST AD over all labs was 10.29 (range: 8.38-12.94) without the ONT-cgMLST-Polisher and 1.26 (range: 0.56-1.81) with the ONT-cgMLST-Polisher (Table 4a). The mean average number of missing cgMLST targets increased from 24.11 to 39.85 when the ONT-cgMLST-Polisher was applied, while the GT itself had an average of 14.31 missing targets. Notably, not all samples were equally affected. Of the 16, only three to six samples had a distance to the GT at all, but a quite high maximum AD of 49 to 75 without and 4 to 14 with the ONT-cgMLST-Polisher. Overall, the errors were either species-specific, e.g., the *E. faecium* samples had hardly any inaccuracy, or strain-specific for the other three species distributed (Fig. S1-S4).

**TABLE 4.** Error analysis results of the 16 ring trial isolates stratified by participating laboratories. Basecalling was performed with Dorado SUP m4.3 (4a) and Dorado SUP m5.0 (4b), respectively. All datasets were analyzed with and without the ONT-cgMLST-Polisher. Distances were calculated against the GT.

| (a) |  | Samples with Distance to GT | Max. cgMLST Allele Distance | Avg. cgMLST Allele Distance (SD) | Avg. Missing cgMLST Alleles (SD) | Max. Missing cgMLST Alleles | Avg. Ns Called | Avg. Coverage Assembled |
| --- | --- | --- | --- | --- | --- | --- | --- | --- |
|  | GT |  |  |  | **14.31 (16.00)** | **71** |  |  |
|  | Lab 1 | 6 | 75 | 12.94 (24.58) | 29.00 (28.25) | 90 | n.a.* | 78 |
|  | Lab 2 | 5 | 68 | 11.44 (21.69) | 25.06 (22.42) | 71 | n.a. | 155 |
|  | Lab 3 | 5 | 49 | 9.00 (17.09) | 22.75 (19.60) | 71 | n.a. | 266 |
|  | Lab 4 | 4 | 61 | 8.38 (17.25) | 19.75 (17.04) | 71 | n.a. | 233 |
|  | Lab 5 | 4 | 46 | 8.38 (16.32) | 20.38 (17.01) | 71 | n.a. | 198 |
|  | Lab 6 | 5 | 69 | 11.63 (21.83) | 27.75 (26.65) | 87 | n.a. | 215 |
|  | Average |  |  | **10.29** | **24.11 (21.83)** |  |  |  |
| ONT-cgMLST-Polisher | Lab 1 | 4 | 11 | 1.69 (3.44) | 47.88 (55.38) | 155 | 53.63 | 78 |
|  | Lab 2 | 4 | 14 | 1.63 (3.79) | 41.81 (46.19) | 126 | 46.63 | 155 |
|  | Lab 3 | 3 | 10 | 1.13 (2.73) | 36.94 (37.98) | 98 | 36.75 | 266 |
|  | Lab 4 | 4 | 4 | 0.56 (1.21) | 32.38 (31.78) | 98 | 31.06 | 233 |
|  | Lab 5 | 3 | 6 | 0.75 (1.73) | 33.00 (31.79) | 83 | 32.69 | 198 |
|  | Lab 6 | 3 | 13 | 1.81 (4.05) | 47.13 (55.54) | 158 | 51.38 | 215 |
|  | Average |  |  | **1.26** | **39.85 (43.11)** |  | **42.02** |  |
| (b) |  | Samples with Distance to GT | Max. cgMLST Allele Distance | Avg. cgMLST Allele Distance (SD) | Avg. Missing cgMLST Alleles (SD) | Max. Missing cgMLST Alleles | Avg. Ns Called | Avg. Coverage Assembled |
|  | GT |  |  |  | **14.31 (16.00)** | **71** |  |  |
|  | Lab 1 | 4 | 4 | 0.63 (1.26) | 16.25 (15.55) | 71 | n.a. | 74 |
|  | Lab 2 | 4 | 2 | 0.38 (0.72) | 15.38 (15.68) | 71 | n.a. | 155 |
|  | Lab 3 | 2 | 2 | 0.25 (0.68) | 14.81 (15.86) | 71 | n.a. | 276 |
|  | Lab 4 | 4 | 2 | 0.31 (0.60) | 14.69 (15.85) | 71 | n.a. | 243 |
|  | Lab 5 | 3 | 2 | 0.25 (0.58) | 14.69 (15.87) | 71 | n.a. | 206 |
|  | Lab 6 | 5 | 2 | 0.38 (0.62) | 16.31 (15.62) | 71 | n.a. | 224 |
|  | Average |  |  | **0.36** | **15.21 (15.77)** |  |  |  |
| ONT-cgMLST-Polisher | Lab 1 | 3 | 2 | 0.38 (0.81) | 19.56 (15.57) | 71 | 7.88 | 74 |
|  | Lab 2 | 2 | 2 | 0.19 (0.54) | 17.88 (15.53) | 71 | 6.69 | 155 |
|  | Lab 3 | 2 | 2 | 0.25 (0.68) | 15.94 (15.50) | 71 | 3.81 | 276 |
|  | Lab 4 | 1 | 1 | 0.06 (0.25) | 15.69 (15.48) | 71 | 3.06 | 243 |
|  | Lab 5 | 1 | 1 | 0.06 (0.25) | 15.5 (15.61) | 71 | 3.63 | 206 |
|  | Lab 6 | 1 | 1 | 0.06 (0.25) | 20.44 (16.69) | 71 | 13.31 | 224 |
|  | Average |  |  | **0.17** | **17.17 (15.74)** |  | **6.40** |  |

\* n.a. – not applicable

For data basecalled with Dorado SUP m5.0, the results improved substantially (Table 4b). Here, the mean average AD was 0.36 without and 0.17 with the ONT-cgMLST-Polisher and the mean average number of missing data reduced to 15.21 and 17.17 without and with the ONT-cgMLST-Polisher, respectively. This is in line with the average number of Ns called by the ONT-cgMLST-Polisher, which decreased from 42.02 to 6.40. The maximum cgMLST AD in a sample was reduced from four to one allele. Most recently, we tested the novel (January 2025) Dorado release v0.9.1, for which a significant basecalling speed improvement is claimed (https://github.com/nanoporetech/dorado/releases), using the ring trial dataset of laboratory 4 with 17.02 Gb basecalled bases (Table 3). Indeed, we could detect a significant speed improvement for model 5.0 basecalling with 18.7 hours (version 0.9.1) versus 46.3 hours (version 0.8.3) for re-basecalling (Table S3).

## Discussion

In this study, we first used a set of 80 MDRO to test the most recent available bioinformatics tools for ONT data processing. We found that Medaka v2.0 with the bacterial methylation model results in a substantial error reduction in comparison to Medaka v1.12 with both Dorado SUP m4.3 and m5.0 data. Still, the error rate for data basecalled with Dorado SUP m4.3 is occasionally too high, while data basecalled with Dorado SUP m5.0 had almost no errors in our samples. In line with this, the ONT-cgMLST-Polisher improved Dorado SUP m4.3 data substantially, for the price of a slightly higher number of missing targets, whereas no effect on the error-rate was noted with Dorado SUP m5.0 data. We are not the first claiming to have achieved accurate bacterial genomic genotyping results from ONT data with a tertiary correction. Chiou *et al.* developed a quite compute-intensive reference-based-method, Modpolish, for correcting ONT base-modification-mediated errors (19). When evaluating this tool with some of our data, we got sometimes polished sequences with (nearly) no errors but often sequences with newly introduced errors. The authors of a more recent study used a deep learning variant caller, Clair3, to process ONT data from 14 diverse bacterial species for SNP and indel variant calling (20, 21). When reanalyzing the 14 ONT raw datasets, it turned out that they showed hardly any errors caused by methylation. Finally, Fuchs *et al.* again used - among others - Clair3 in their new tool, NanoCore, to calculate and visualize cgMLST-like sample distances (22). Also, this tool is quite compute-intensive and produced not always sufficient results with our data (data not shown). With respect to the recovery of chromosomal or plasmid-borne location of AMR targets, ONT was superior to Illumina data (Table 2). Interestingly ONT data called with Dorado SUP m4.3 performed better than with Dorado SUP m5.0. Here, however, not the accuracy but the contiguity of assemblies was evaluated.

To assess the reproducibility of our findings, we organized a ring trial with six laboratories that performed ONT sequencing on the 16 isolates including some of the most challenging samples of the 80 MDRO isolates. Again, we found that data basecalled with Dorado SUP m4.3 occasionally contained too many errors even after polishing with Medaka v2.0 and the ONT-cgMLST-Polisher. Whereas the 16 samples chosen from the evaluation dataset exhibited an average AD to the GT of 0.375 using Dorado SUP m4.3, higher averages were observed with quite some variation between the laboratories (Table 4a). For data basecalled with Dorado SUP m5.0 and polished with Medaka v2.0, assemblies could be further improved by applying the ONT-cgMLST-Polisher and resulted in nearly perfect data (Table 4b) with only marginally higher numbers of missing targets. Overall, using the ONT-cgMLST-Polisher safeguards for the possibility of methylation-patterns that the Dorado and Medaka models were not trained for.

In addition to testing the reproducibility of post-processing tools, the ring trial provided some insights into raw data acquisition. Neither the total number nor the sequencing number of pores at the start are reliable indicators for final data output in terms of bases called. However, the total number and/or sequencing number of pores after 90 minutes seem to be such a predictor. Further, applying genome size compensation seems to mitigate the effects of uneven average depth of coverage.

We have shown that with the newest polishing tools cgMLST-based analysis of ONT data is now comparable to Illumina sequencing with respect to the error-rate. Moreover, ONT outperforms short-read data in terms of AMR target recovery and correct chromosomal or plasmid-borne localization, which might affect infection control measures in the hospital setting. In January 2025, ONT released Dorado v0.9.1 that shows a significant speed improvement of about 60% for model 5.0 basecalling with newer graphical processing units (GPU; Table S3). Thereby, live basecalling in SUP mode becomes possible with recent GPUs like the Nvidia RTX 4090. For such a ‘gamer’ desktop computer an investment of about 5,000 € is required. Therefore, it would be very desirable that ONT includes Dorado v0.9.1 or newer SUP m5 in MinKNOW soon and ‘freezes’ the R10.4.1 flowcell and at least RBK chemistry for CLIA/ISO accredited laboratories.

Our study has some limitations. First, we sent only DNA instead of living organisms to the six laboratories that participated in the ring trial, and thereby did not test the influence of DNA extraction methods. Second, the accuracy was only determined for cgMLST targets. In accordance with recent practices in public health and clinical microbiology, the intergenic regions were not controlled here, although they are also corrected by the ONT-cgMLST-Polisher. Third, the base patterns of the nucleotide modifications were not analyzed in detail. However, this was done recently by some of the co-authors and there is no reason to believe that something has changed since then (7, 8). Finally, a proprietary tertiary correction tool was evaluated. However, just the improvements of the primary (basecaller) and secondary (Medaka) correction would warrant an own publication.

In conclusion, Dorado SUP 4.3 and even more Dorado SUP 5.0 model combined with Medaka v2.0 and the ONT-cgMLST-Polisher improve ONT sequencing accuracy substantially and make it finally accurate and sufficiently reproducible for whole-genome genotyping based surveillance of bacteria. The rather low capital investment and per-sample consumable costs provide opportunities also for smaller 374 laboratories to perform WGS-based surveillance on a routine basis.

## Authors contributions

Karola Prior and Alexander Mellmann conceptualized the study, analyzed the results, and drafted the manuscript. All authors contributed to the laboratory ring trial work and contributed to the correction of the draft and agreed with its content.

