## Supplementary material for "Accurate and Reproducible Whole-Genome Genotyping for Bacterial Genomic Surveillance with Nanopore Sequencing Data": SupplMaterial

**Supplemental Table 1** Isolates of evaluation dataset (n=80) with cgMLST allelic distances (AD) to ground truth. In **bold** selected 16 isolates for ring trial.

| **Sample ID** | **Species** | **Illumina AD** | **Dorado SUP m4.3 AD** | | | | | | **Dorado SUP m5.0 AD** | | | | | |
| --- | --- | --- | --- | --- | --- | --- | --- | --- | --- | --- | --- | --- | --- | --- |
|  |  |  | **Flye only** | **Medaka v1.12 SUP m4.3** | **Medaka v2.0 bacterial methylation model** | **Flye only** | **Medaka v1.12 SUP m4.3** | **Medaka v2.0 bacterial methylation model** | **Flye only** | **Medaka v1.12 SUP m4.3** | **Medaka v2.0 bacterial methylation model** | **Flye only** | **Medaka v1.12 SUP m4.3** | **Medaka v2.0 bacterial methylation model** |
|  |  |  |  |  |  | **+ ONT-cgMLST-Polisher** | | |  |  |  | **+ ONT-cgMLST-Polisher** | | |
| 24025 | *S. marcescens* | 0 | 15 | 26 | 1 | 0 | 0 | 0 | 10 | 6 | 0 | 1 | 0 | 0 |
| 24100 | *P. aeruginosa* | 0 | 1 | 0 | 0 | 0 | 0 | 0 | 1 | 6 | 0 | 0 | 0 | 0 |
| 24110 | *S. aureus* | 0 | 0 | 0 | 0 | 0 | 0 | 0 | 0 | 0 | 0 | 0 | 0 | 0 |
| 24111 | *S. aureus* | 0 | 1 | 1 | 0 | 0 | 0 | 0 | 0 | 0 | 0 | 0 | 0 | 0 |
| 24212 | *C. freundii* | 0 | 0 | 0 | 0 | 0 | 0 | 0 | 0 | 0 | 0 | 0 | 0 | 0 |
| A24358 | *E. hormaechei* | 0 | 4 | 4 | 2 | 0 | 0 | 0 | 0 | 1 | 0 | 0 | 0 | 0 |
| A24592 | *P. aeruginosa* | 0 | 0 | 0 | 0 | 0 | 0 | 0 | 0 | 0 | 0 | 0 | 0 | 0 |
| A24981 | *E. coli* | 0 | 0 | 0 | 0 | 0 | 0 | 0 | 0 | 0 | 0 | 0 | 0 | 0 |
| A24983 | *E. coli* | 0 | 1 | 1 | 1 | 0 | 0 | 0 | 2 | 0 | 0 | 0 | 0 | 0 |
| **A24994** | ***E. faecium*** | **0** | **0** | **0** | **0** | **0** | **0** | **0** | **0** | **0** | **0** | **0** | **0** | **0** |
| A25355 | *P. mirabilis* | 0 | 0 | 0 | 0 | 0 | 0 | 0 | 0 | 0 | 0 | 0 | 0 | 0 |
| A25726 | *S. marcescens* | 0 | 1 | 2 | 1 | 0 | 0 | 0 | 0 | 0 | 0 | 0 | 0 | 0 |
| A25786 | *S. aureus* | 0 | 0 | 0 | 0 | 0 | 0 | 0 | 0 | 0 | 0 | 0 | 0 | 0 |
| A25934 | *S. aureus* | 0 | 0 | 0 | 0 | 0 | 0 | 0 | 0 | 0 | 0 | 0 | 0 | 0 |
| A25962 | *E. faecium* | 0 | 0 | 0 | 0 | 0 | 0 | 0 | 0 | 0 | 0 | 0 | 0 | 0 |
| A26107 | *K. pneumoniae* | 0 | 1 | 0 | 0 | 0 | 0 | 0 | 0 | 1 | 0 | 0 | 0 | 0 |
| **A26322** | ***E. hormaechei*** | **0** | **19** | **17** | **2** | **0** | **0** | **0** | **10** | **4** | **0** | **0** | **0** | **0** |
| A26326 | *A. baumannii* | 0 | 0 | 0 | 0 | 0 | 0 | 0 | 2 | 1 | 0 | 0 | 0 | 0 |
| A26329 | *A. baumannii* | 0 | 4 | 4 | 1 | 0 | 0 | 0 | 1 | 2 | 0 | 0 | 0 | 0 |
| A26335 | *E. faecium* | 0 | 0 | 0 | 0 | 0 | 0 | 0 | 0 | 0 | 0 | 0 | 0 | 0 |
| A26362 | *E. hormaechei* | 0 | 0 | 0 | 0 | 0 | 0 | 0 | 0 | 0 | 0 | 0 | 0 | 0 |
| **A26371** | ***E. hormaechei*** | **0** | **74** | **71** | **41** | **4** | **4** | **3** | **1** | **2** | **1** | **1** | **1** | **1** |
| **A26406** | ***K. pneumoniae*** | **0** | **0** | **0** | **0** | **0** | **0** | **0** | **0** | **0** | **0** | **0** | **0** | **0** |
| A26461 | *P. aeruginosa* | 0 | 0 | 0 | 0 | 0 | 0 | 0 | 0 | 0 | 0 | 0 | 0 | 0 |
| **A26462** | ***M. morganii*** | **0** | **0** | **0** | **0** | **0** | **0** | **0** | **0** | **0** | **0** | **0** | **0** | **0** |
| A26535 | *E. hormaechei* | 0 | 1 | 1 | 0 | 0 | 0 | 0 | 0 | 1 | 0 | 0 | 0 | 0 |
| A26715 | *S. marcescens* | 0 | 0 | 0 | 0 | 0 | 0 | 0 | 0 | 0 | 0 | 0 | 0 | 0 |
| A26716 | *S. marcescens* | 0 | 9 | 8 | 0 | 0 | 0 | 0 | 3 | 1 | 0 | 0 | 0 | 0 |
| **A26728** | ***E. faecium*** | **0** | **0** | **0** | **0** | **0** | **0** | **0** | **0** | **0** | **0** | **0** | **0** | **0** |
| A27084 | *K. pneumoniae* | 0 | 0 | 0 | 0 | 0 | 0 | 0 | 0 | 0 | 0 | 0 | 0 | 0 |
| A27088 | *E. coli* | 0 | 1 | 0 | 0 | 0 | 0 | 0 | 1 | 2 | 0 | 0 | 0 | 0 |
| A27312 | *P. aeruginosa* | 0 | 0 | 0 | 0 | 0 | 0 | 0 | 0 | 0 | 0 | 0 | 0 | 0 |
| A27468 | *E. coli* | 0 | 0 | 0 | 0 | 0 | 0 | 0 | 0 | 0 | 0 | 0 | 0 | 0 |
| A27543 | *S. aureus* | 0 | 1 | 0 | 0 | 0 | 0 | 0 | 0 | 0 | 0 | 0 | 0 | 0 |
| A27547 | *C. freundii* | 0 | 0 | 0 | 0 | 0 | 0 | 0 | 0 | 0 | 0 | 0 | 0 | 0 |
| A27667 | *E. coli* | 0 | 0 | 1 | 0 | 0 | 0 | 0 | 0 | 0 | 0 | 0 | 0 | 0 |
| A27672 | *P. mirabilis* | 0 | 0 | 0 | 0 | 0 | 0 | 0 | 0 | 0 | 0 | 0 | 0 | 0 |
| **A27739** | ***K. pneumoniae*** | **0** | **2** | **3** | **0** | **0** | **0** | **0** | **0** | **0** | **0** | **0** | **0** | **0** |
| A27745 | *M. morganii* | 0 | 0 | 0 | 0 | 0 | 0 | 0 | 0 | 0 | 0 | 0 | 0 | 0 |
| A27776 | *E. hormaechei* | 0 | 1 | 3 | 1 | 0 | 0 | 0 | 0 | 0 | 0 | 0 | 0 | 0 |
| A27780 | *K. pneumoniae* | 0 | 0 | 0 | 0 | 0 | 0 | 0 | 0 | 0 | 0 | 0 | 0 | 0 |
| A27855 | *E. coli* | 0 | 1 | 2 | 0 | 0 | 0 | 0 | 0 | 0 | 0 | 0 | 0 | 0 |
| A27857 | *K. pneumoniae* | 0 | 0 | 0 | 0 | 0 | 0 | 0 | 0 | 0 | 0 | 0 | 0 | 0 |
| **A28064** | ***K. pneumoniae*** | **1** | **98** | **98** | **57** | **2** | **2** | **1** | **0** | **0** | **0** | **0** | **0** | **0** |
| A28160 | *E. coli* | 0 | 1 | 2 | 0 | 0 | 0 | 0 | 0 | 2 | 0 | 0 | 0 | 0 |
| A28161 | *K. pneumoniae* | 0 | 2 | 0 | 0 | 0 | 0 | 0 | 0 | 4 | 0 | 0 | 0 | 0 |
| A28164 | *E. coli* | 0 | 0 | 0 | 0 | 0 | 0 | 0 | 0 | 0 | 0 | 0 | 0 | 0 |
| A28340 | *E. coli* | 0 | 7 | 4 | 0 | 0 | 0 | 0 | 3 | 1 | 0 | 0 | 0 | 0 |
| A28573 | *S. aureus* | 0 | 0 | 0 | 0 | 0 | 0 | 0 | 0 | 0 | 0 | 0 | 0 | 0 |
| A28602 | *P. aeruginosa* | 0 | 0 | 0 | 0 | 0 | 0 | 0 | 1 | 0 | 0 | 1 | 0 | 0 |
| **A28603** | ***M. morganii*** | **0** | **0** | **0** | **0** | **0** | **0** | **0** | **0** | **1** | **0** | **0** | **0** | **0** |
| A28879 | *S. aureus* | 0 | 0 | 0 | 0 | 0 | 0 | 0 | 0 | 0 | 0 | 0 | 0 | 0 |
| A28999 | *S. marcescens* | 0 | 0 | 0 | 0 | 0 | 0 | 0 | 0 | 0 | 0 | 0 | 0 | 0 |
| A29003 | *P. mirabilis* | 0 | 1 | 2 | 1 | 1 | 2 | 1 | 1 | 1 | 1 | 1 | 1 | 1 |
| **A29005** | ***M. morganii*** | **0** | **33** | **38** | **21** | **1** | **1** | **1** | **0** | **0** | **0** | **0** | **0** | **0** |
| A29006 | *P. mirabilis* | 0 | 1 | 1 | 0 | 0 | 0 | 0 | 0 | 0 | 0 | 0 | 0 | 0 |
| A29171 | *S. aureus* | 0 | 0 | 0 | 0 | 0 | 0 | 0 | 0 | 0 | 0 | 0 | 0 | 0 |
| A29251 | *S. aureus* | 0 | 0 | 0 | 0 | 0 | 0 | 0 | 0 | 0 | 0 | 0 | 0 | 0 |
| A29253 | *S. aureus* | 0 | 3 | 2 | 0 | 0 | 0 | 0 | 0 | 0 | 0 | 0 | 0 | 0 |
| A29301 | *S. aureus* | 0 | 3 | 1 | 0 | 0 | 0 | 0 | 1 | 0 | 0 | 0 | 0 | 0 |
| A29411 | *K. pneumoniae* | 0 | 1 | 2 | 1 | 0 | 0 | 0 | 1 | 1 | 0 | 0 | 0 | 0 |
| A29484 | *S. aureus* | 0 | 4 | 3 | 0 | 0 | 0 | 0 | 1 | 0 | 0 | 0 | 0 | 0 |
| A29767 | *K. pneumoniae* | 0 | 0 | 0 | 0 | 0 | 0 | 0 | 0 | 0 | 0 | 0 | 0 | 0 |
| **A29816** | ***E. hormaechei*** | **0** | **0** | **0** | **0** | **0** | **0** | **0** | **0** | **0** | **0** | **0** | **0** | **0** |
| A29847 | *P. aeruginosa* | 0 | 4 | 8 | 0 | 1 | 1 | 0 | 3 | 3 | 0 | 0 | 0 | 0 |
| **A29871** | ***E. hormaechei*** | **0** | **3** | **5** | **0** | **0** | **0** | **0** | **4** | **2** | **0** | **0** | **0** | **0** |
| A30424 | *E. hormaechei* | 0 | 0 | 0 | 0 | 0 | 0 | 0 | 0 | 0 | 0 | 0 | 0 | 0 |
| **A30987** | ***K. pneumoniae*** | **0** | **3** | **3** | **0** | **0** | **0** | **0** | **0** | **0** | **0** | **0** | **0** | **0** |
| A31005 | *P. aeruginosa* | 0 | 11 | 14 | 0 | 0 | 0 | 0 | 8 | 6 | 0 | 0 | 0 | 0 |
| **A31008** | ***E. faecium*** | **0** | **0** | **0** | **0** | **0** | **0** | **0** | **0** | **0** | **0** | **0** | **0** | **0** |
| A31034 | *P. mirabilis* | 0 | 1 | 2 | 0 | 0 | 1 | 0 | 0 | 0 | 0 | 0 | 0 | 0 |
| A31131 | *S. marcescens* | 0 | 0 | 0 | 0 | 0 | 0 | 0 | 0 | 0 | 0 | 0 | 0 | 0 |
| A31138 | *S. marcescens* | 0 | 0 | 0 | 0 | 0 | 0 | 0 | 0 | 0 | 0 | 0 | 0 | 0 |
| **A31143** | ***M. morganii*** | **0** | **29** | **35** | **12** | **3** | **2** | **1** | **0** | **0** | **0** | **0** | **0** | **0** |
| A31232 | *E. coli* | 0 | 0 | 0 | 0 | 0 | 0 | 0 | 0 | 0 | 0 | 0 | 0 | 0 |
| A31645 | *P. aeruginosa* | 0 | 0 | 0 | 0 | 0 | 0 | 0 | 0 | 0 | 0 | 0 | 0 | 0 |
| A31772 | *P. aeruginosa* | 0 | 1 | 1 | 0 | 0 | 0 | 0 | 0 | 1 | 0 | 0 | 0 | 0 |
| A32326 | *P. aeruginosa* | 0 | 0 | 0 | 0 | 0 | 0 | 0 | 0 | 0 | 0 | 0 | 0 | 0 |
| A32614 | *E. hormaechei* | 0 | 5 | 3 | 0 | 0 | 0 | 0 | 1 | 1 | 1 | 0 | 0 | 0 |
| **A33124** | ***E. faecium*** | **0** | **14** | **27** | **0** | **0** | **0** | **0** | **6** | **8** | **0** | **0** | **0** | **0** |

**Supplemental Table 2** Coverage and cgMLST allelic distance (AD) to ground truth per ring trial isolate and participating laboratory. Medaka v2.0 bacterial methylation model was used for polishing.

| **Sample ID** | **Species** | **Lab number** | **Coverage assembled (m4.3/m5.0)** | **Dorado SUP m4.3 AD** | **Dorado SUP m4.3 + ONT-cgMLST-Polisher AD** | **Dorado SUP m5.0 AD** | **Dorado SUP m5.0 + ONT-cgMLST-Polisher AD** |
| --- | --- | --- | --- | --- | --- | --- | --- |
| A24994 | *E. faecium* | Lab 1 | 117/109 | 0 | 0 | 0 | 0 |
|  |  | Lab 2 | 226/231 | 0 | 0 | 0 | 0 |
|  |  | Lab 3 | 454/476 | 0 | 0 | 0 | 0 |
|  |  | Lab 4 | 255/278 | 0 | 0 | 0 | 0 |
|  |  | Lab 5 | 205/203 | 0 | 0 | 0 | 0 |
|  |  | Lab 6 | 334/332 | 0 | 0 | 0 | 0 |
| A26322 | *E. hormaechei* | Lab 1 | 52/49 | 1 | 0 | 1 | 0 |
|  |  | Lab 2 | 129/128 | 1 | 0 | 0 | 0 |
|  |  | Lab 3 | 190/198 | 0 | 0 | 0 | 0 |
|  |  | Lab 4 | 213/224 | 0 | 0 | 1 | 0 |
|  |  | Lab 5 | 210/216 | 0 | 0 | 0 | 0 |
|  |  | Lab 6 | 140/142 | 0 | 0 | 1 | 0 |
| A26371 | *E. hormaechei* | Lab 1 | 80/76 | 75 | 11 | 4 | 2 |
|  |  | Lab 2 | 132/131 | 68 | 14 | 2 | 2 |
|  |  | Lab 3 | 205/212 | 49 | 10 | 2 | 2 |
|  |  | Lab 4 | 228/237 | 32 | 4 | 1 | 1 |
|  |  | Lab 5 | 129/134 | 46 | 6 | 1 | 1 |
|  |  | Lab 6 | 311/326 | 69 | 13 | 2 | 1 |
| A26404 | *K. pneumoniae* | Lab 1 | 62/59 | 0 | 0 | 0 | 0 |
|  |  | Lab 2 | 78/77 | 0 | 0 | 1 | 0 |
|  |  | Lab 3 | 125/130 | 0 | 0 | 0 | 0 |
|  |  | Lab 4 | 167/174 | 0 | 0 | 0 | 0 |
|  |  | Lab 5 | 119/125 | 0 | 0 | 0 | 0 |
|  |  | Lab 6 | 191/203 | 0 | 0 | 0 | 0 |
| A26462 | *M. morganii* | Lab 1 | 46/44 | 0 | 0 | 0 | 0 |
|  |  | Lab 2 | 84/84 | 0 | 0 | 0 | 0 |
|  |  | Lab 3 | 414/429 | 0 | 0 | 0 | 0 |
|  |  | Lab 4 | 205/214 | 0 | 0 | 0 | 0 |
|  |  | Lab 5 | 153/163 | 0 | 0 | 0 | 0 |
|  |  | Lab 6 | 181/192 | 0 | 0 | 0 | 0 |
| A26728 | *E. faecium* | Lab 1 | 184/169 | 0 | 0 | 0 | 0 |
|  |  | Lab 2 | 270/269 | 0 | 0 | 0 | 0 |
|  |  | Lab 3 | 262/271 | 0 | 0 | 0 | 0 |
|  |  | Lab 4 | 330/346 | 0 | 0 | 0 | 0 |
|  |  | Lab 5 | 158/159 | 0 | 0 | 0 | 0 |
|  |  | Lab 6 | 233/239 | 0 | 0 | 0 | 0 |

| **Sample ID** | **Species** | **Lab number** | **Coverage assembled (m4.3/m5.0)** | **Dorado SUP m4.3 AD** | **Dorado SUP m4.3 + ONT-cgMLST-Polisher AD** | **Dorado SUP m5.0 AD** | **Dorado SUP m5.0 + ONT-cgMLST-Polisher AD** |
| --- | --- | --- | --- | --- | --- | --- | --- |
| A27739 | *K. pneumoniae* | Lab 1 | 63/61 | 0 | 0 | 0 | 0 |
|  |  | Lab 2 | 166/166 | 0 | 0 | 0 | 0 |
|  |  | Lab 3 | 132/139 | 0 | 0 | 0 | 0 |
|  |  | Lab 4 | 199/207 | 0 | 0 | 0 | 0 |
|  |  | Lab 5 | 113/115 | 0 | 0 | 0 | 0 |
|  |  | Lab 6 | 111/114 | 0 | 0 | 0 | 0 |
| A228064 | *K. pneumoniae* | Lab 1 | 44/42 | 61 | 3 | 0 | 0 |
|  |  | Lab 2 | 58/57 | 49 | 1 | 0 | 0 |
|  |  | Lab 3 | 111/115 | 46 | 0 | 0 | 0 |
|  |  | Lab 4 | 146/155 | 61 | 1 | 0 | 0 |
|  |  | Lab 5 | 143/148 | 46 | 0 | 0 | 0 |
|  |  | Lab 6 | 232/245 | 35 | 0 | 0 | 0 |
| A28603 | *M. morganii* | Lab 1 | 62/60 | 0 | 0 | 0 | 0 |
|  |  | Lab 2 | 129/127 | 0 | 0 | 0 | 0 |
|  |  | Lab 3 | 386/398 | 0 | 0 | 0 | 0 |
|  |  | Lab 4 | 378/390 | 0 | 0 | 0 | 0 |
|  |  | Lab 5 | 244/249 | 0 | 0 | 0 | 0 |
|  |  | Lab 6 | 260/274 | 0 | 0 | 1 | 0 |
| A29005 | *M. morganii* | Lab 1 | 85/81 | 37 | 7 | 2 | 2 |
|  |  | Lab 2 | 92/92 | 35 | 5 | 1 | 1 |
|  |  | Lab 3 | 308/313 | 26 | 4 | 2 | 2 |
|  |  | Lab 4 | 126/130 | 21 | 1 | 0 | 0 |
|  |  | Lab 5 | 291/304 | 22 | 3 | 1 | 0 |
|  |  | Lab 6 | 221/233 | 36 | 7 | 1 | 0 |
| A29816 | *E. hormaechei* | Lab 1 | 35/34 | 0 | 0 | 0 | 0 |
|  |  | Lab 2 | 265/263 | 0 | 0 | 0 | 0 |
|  |  | Lab 3 | 131/136 | 0 | 0 | 0 | 0 |
|  |  | Lab 4 | 347/354 | 0 | 0 | 0 | 0 |
|  |  | Lab 5 | 222/231 | 0 | 0 | 0 | 0 |
|  |  | Lab 6 | 342/361 | 0 | 0 | 0 | 0 |
| A29871 | *E. hormaechei* | Lab 1 | 51/48 | 0 | 0 | 0 | 0 |
|  |  | Lab 2 | 143/141 | 0 | 0 | 0 | 0 |
|  |  | Lab 3 | 300/314 | 0 | 0 | 0 | 0 |
|  |  | Lab 4 | 79/83 | 0 | 0 | 0 | 0 |
|  |  | Lab 5 | 289/295 | 0 | 0 | 0 | 0 |
|  |  | Lab 6 | 204/207 | 0 | 0 | 0 | 0 |
| A30987 | *K. pneumoniae* | Lab 1 | 45/43 | 0 | 0 | 0 | 0 |
|  |  | Lab 2 | 64/64 | 0 | 0 | 0 | 0 |
|  |  | Lab 3 | 89/83 | 1 | 0 | 0 | 0 |
|  |  | Lab 4 | 158/162 | 0 | 0 | 2 | 0 |
|  |  | Lab 5 | 105/109 | 0 | 0 | 0 | 0 |
|  |  | Lab 6 | 103/108 | 1 | 0 | 0 | 0 |
| **Sample ID** | **Species** | **Lab number** | **Coverage assembled (m4.3/m5.0)** | **Dorado SUP m4.3 AD** | **Dorado SUP m4.3 + ONT-cgMLST-Polisher AD** | **Dorado SUP m5.0 AD** | **Dorado SUP m5.0 + ONT-cgMLST-Polisher AD** |
| A31008 | *E. faecium* | Lab 1 | 71/67 | 0 | 0 | 0 | 0 |
|  |  | Lab 2 | 166/165 | 0 | 0 | 0 | 0 |
|  |  | Lab 3 | 427/437 | 0 | 0 | 0 | 0 |
|  |  | Lab 4 | 376/392 | 0 | 0 | 0 | 0 |
|  |  | Lab 5 | 364/388 | 0 | 0 | 0 | 0 |
|  |  | Lab 6 | 250/269 | 0 | 0 | 0 | 0 |
| A31143 | *M. morganii* | Lab 1 | 131/128 | 32 | 7 | 3 | 2 |
|  |  | Lab 2 | 191/190 | 30 | 6 | 2 | 0 |
|  |  | Lab 3 | 209/218 | 22 | 0 | 4 | 0 |
|  |  | Lab 4 | 106/110 | 20 | 3 | 1 | 0 |
|  |  | Lab 5 | 281/291 | 20 | 3 | 2 | 0 |
|  |  | Lab 6 | 132/139 | 45 | 9 | 1 | 0 |
| A33124 | *E. faecium* | Lab 1 | 119/115 | 1 | 0 | 0 | 0 |
|  |  | Lab 2 | 290/297 | 0 | 0 | 0 | 0 |
|  |  | Lab 3 | 520/544 | 0 | 0 | 0 | 0 |
|  |  | Lab 4 | 422/433 | 0 | 0 | 0 | 0 |
|  |  | Lab 5 | 143/166 | 0 | 0 | 0 | 0 |
|  |  | Lab 6 | 197/207 | 0 | 0 | 0 | 0 |

**Supplemental Table 3** Wall-clock time (hours) of two Dorado versions using POD5 files of a RBK run with an output of 17.02 Gb called bases. SUP re-basecalling was done with a Lenovo Legion Pro7i Gen9 laptop with a GeoForce RTX 4090 16GB GDDR6 (Ada Lovelace with Compute Capability 8.9) GPU.

| **Dorado Version** | **Model** | **Hours** | **Factor** |
| --- | --- | --- | --- |
| 0.8.3 | 4.3.0 | 18.4 | 1x |
| 0.8.3 | 5.0.0 | 46.3 | 2.52x |
| 0.9.1 | 5.0.0 | 18.7 | 1.02x |

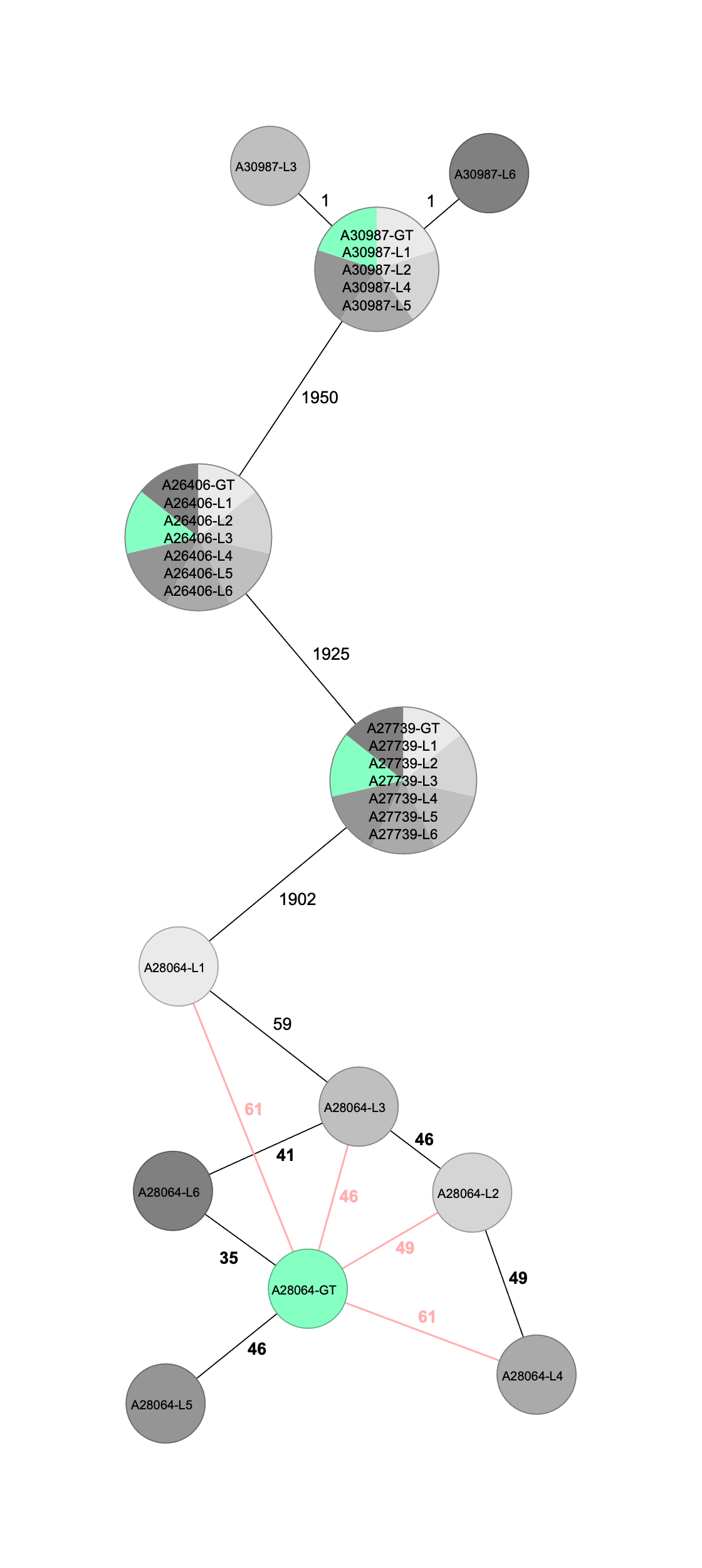
**(a)**

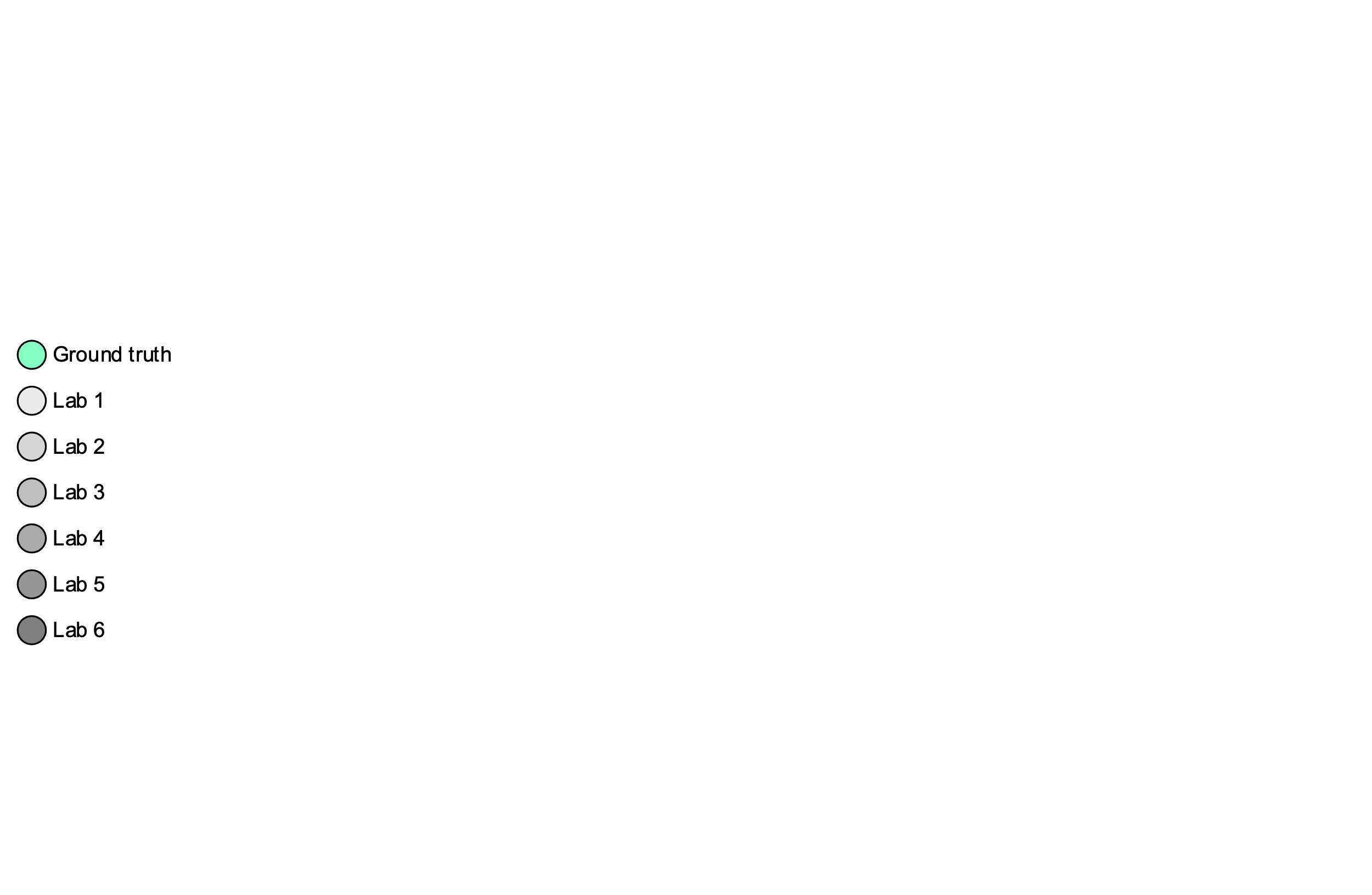

| **(b)** | **Avg. No. of Samples with Distance to GT** | **Max. cgMLST Allele Distance** | **Avg. cgMLST Allele Distance (SD)** | **Max. Missing cgMLST Alleles** | **Avg. Missing cgMLST Alleles (SD)** | **Avg. Ns Called** |
| --- | --- | --- | --- | --- | --- | --- |
| GT |  |  |  | 15 | 9.75 (3.59) |  |
| SUP m4.3 | 1.33 | 61 | 12.50 (24.78) | 41 | 16.29 (11.99) | n.a.* |
| SUP m4.3 + Polisher | 0.50 | 2 | 0.17 (0.33) | 108 | 32.17 (38.64) | 51.29 |
| SUP m5.0 | 0.00 | 2 | 0.13 (0.25) | 18 | 10.38 (3.85) | n.a. |
| SUP m5.0 + Polisher | 0.00 | 0 | 0.00 (0.00) | 22 | 12.25 (3.60) | 7.50 |
| * n.a. – not applicable | | | | | | |

**Supplemental Figure 1**	(a) Minimum spanning tree of *K. pneumoniae* (SUP m4.3, Medaka 2.0) cgMLST data without ONT cgMLST Polisher. Distances are based on cgMLST scheme of *K. pneumoniae* sensu lato (2,358 genes), pairwise ignoring missing values. The values on the connecting lines indicate the number of allelic distances between the connected isolates. Hybracter hybrid assemblies used as ground truth (green). (b) Error analysis results averaged over all sequenced ring trial isolates (n=96) with Dorado basecalling models SUP 4.3 and 5.0 with and without ONT cgMLST Polisher analysed.

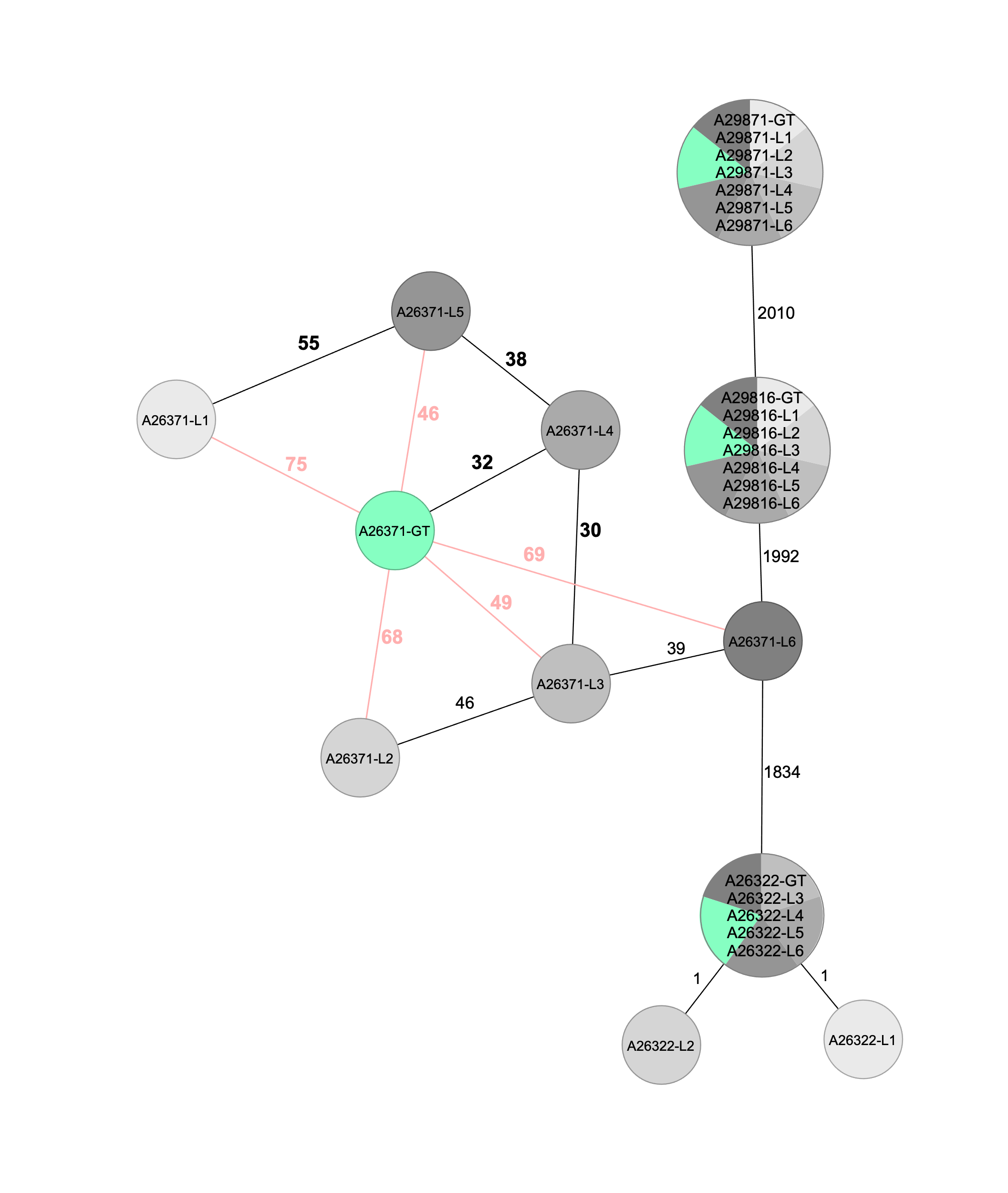
**(a)**

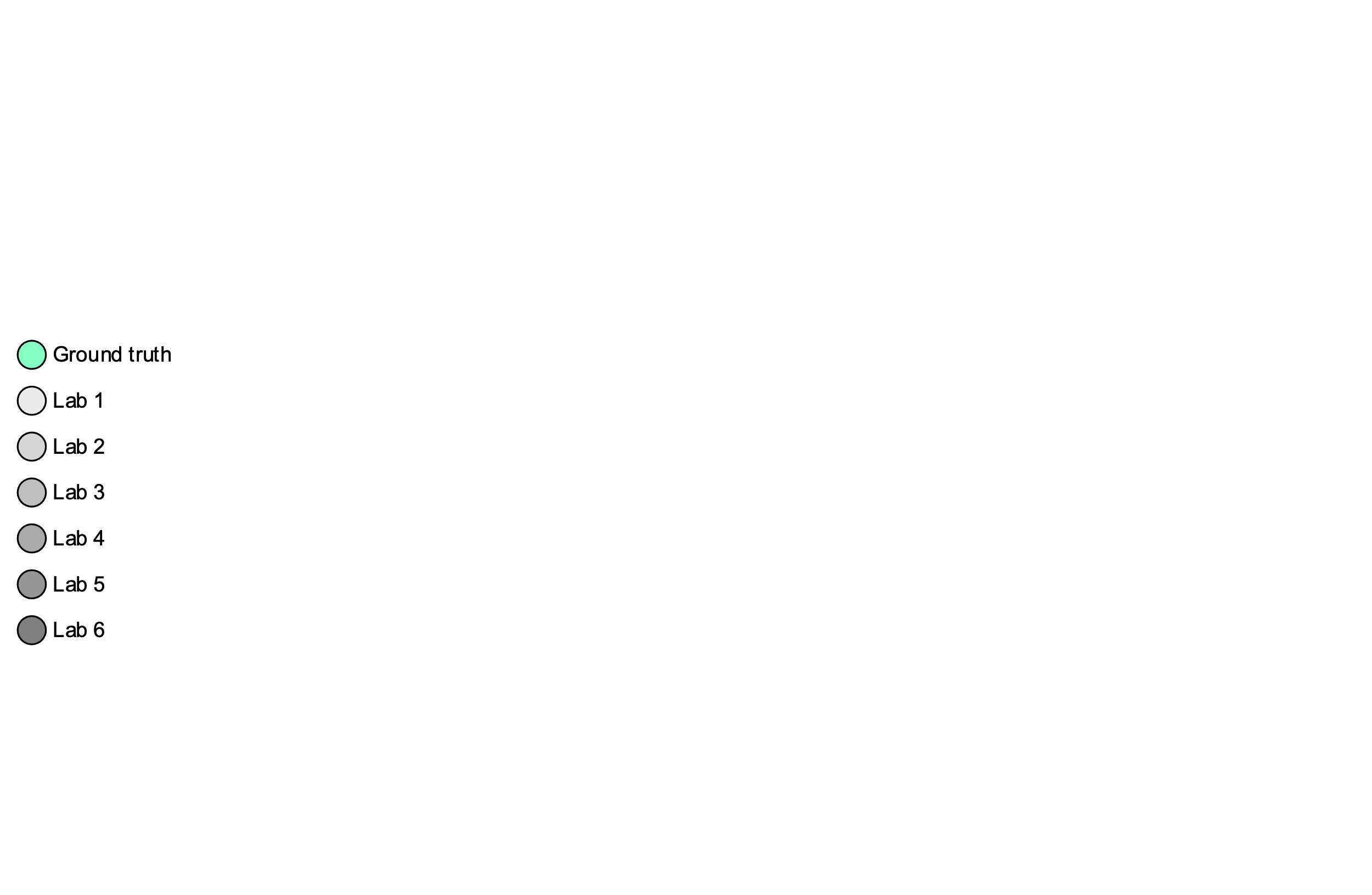

| **(b)** | **Avg. No. of Samples with Distance to GT** | **Max. cgMLST Allele Distance** | **Avg. cgMLST Allele Distance (SD)** | **Max. Missing cgMLST Alleles** | **Avg. Missing cgMLST Alleles (SD)** | **Avg. Ns Called** |
| --- | --- | --- | --- | --- | --- | --- |
| GT |  |  |  | 71 | 25.75 (30.41) |  |
| SUP m4.3 | 1.33 | 75 | 14.21 (28.20) | 90 | 40.00 (33.42) | n.a.* |
| SUP m4.3 + Polisher | 1 | 14 | 2.42 (4.83) | 155 | 53.46 (48.31) | 50.67 |
| SUP m5.0 | 1.5 | 4 | 0.63 (0.99) | 71 | 27.00 (29.58) | n.a. |
| SUP m5.0 + Polisher | 1 | 2 | 0.38 (0.75) | 71 | 28.71 (28.27) | 6.79 |
| * n.a. – not applicable | | | | | | |

**Supplemental Figure 2** (a) Minimum spanning tree of *E. hormaechei* (SUP m4.3, Medaka 2.0) cgMLST data without ONT cgMLST Polisher. Distances are based on cgMLST scheme of *E. hormaechei* (2,178 genes), pairwise ignoring missing values. The values on the connecting lines indicate the number of allelic distances between the connected isolates. Hybracter hybrid assemblies used as ground truth (green). (b) Error analysis results averaged over all sequenced ring trial isolates (n=96) with Dorado basecalling models SUP 4.3 and 5.0 with and without ONT cgMLST Polisher analysed.

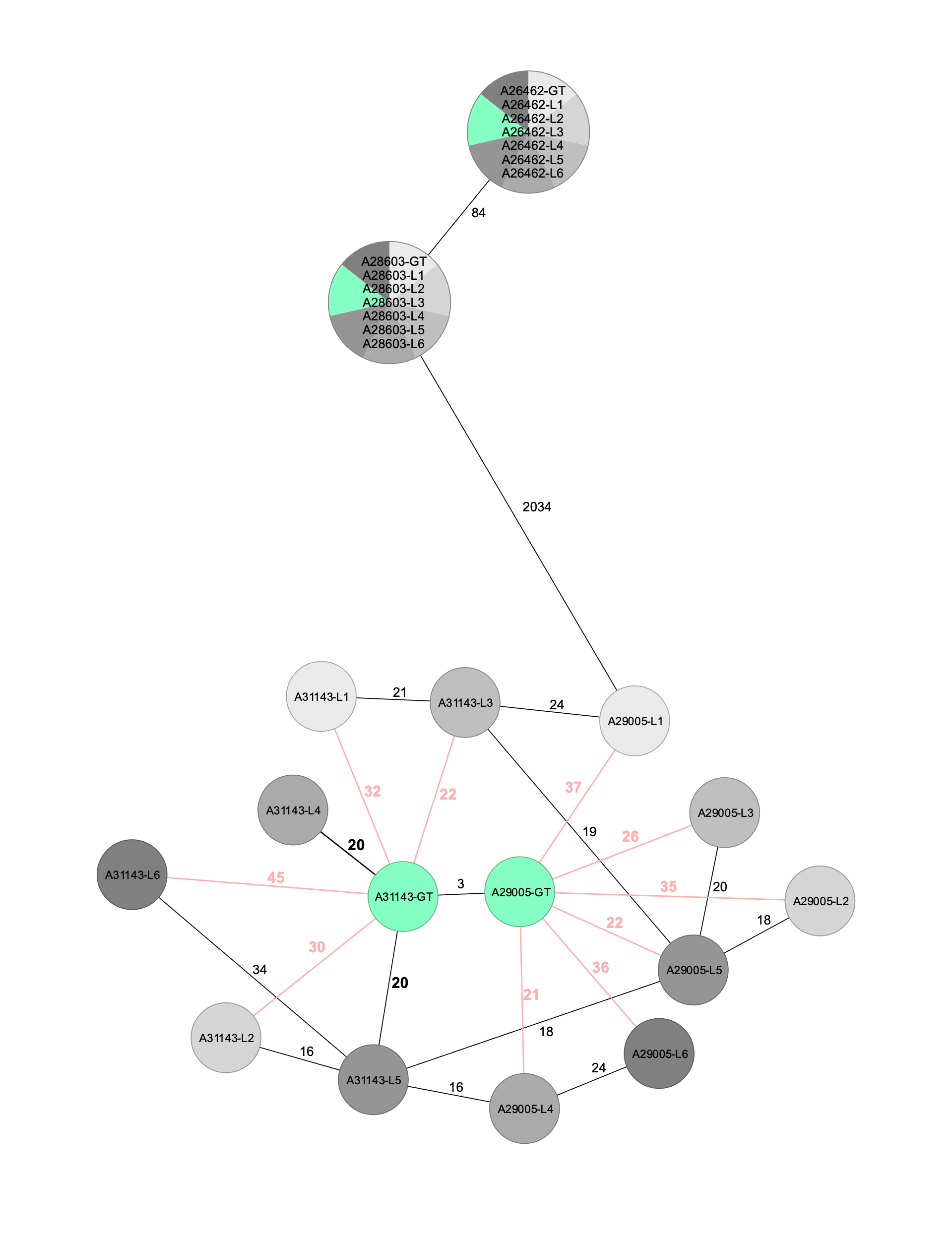
**(a)**

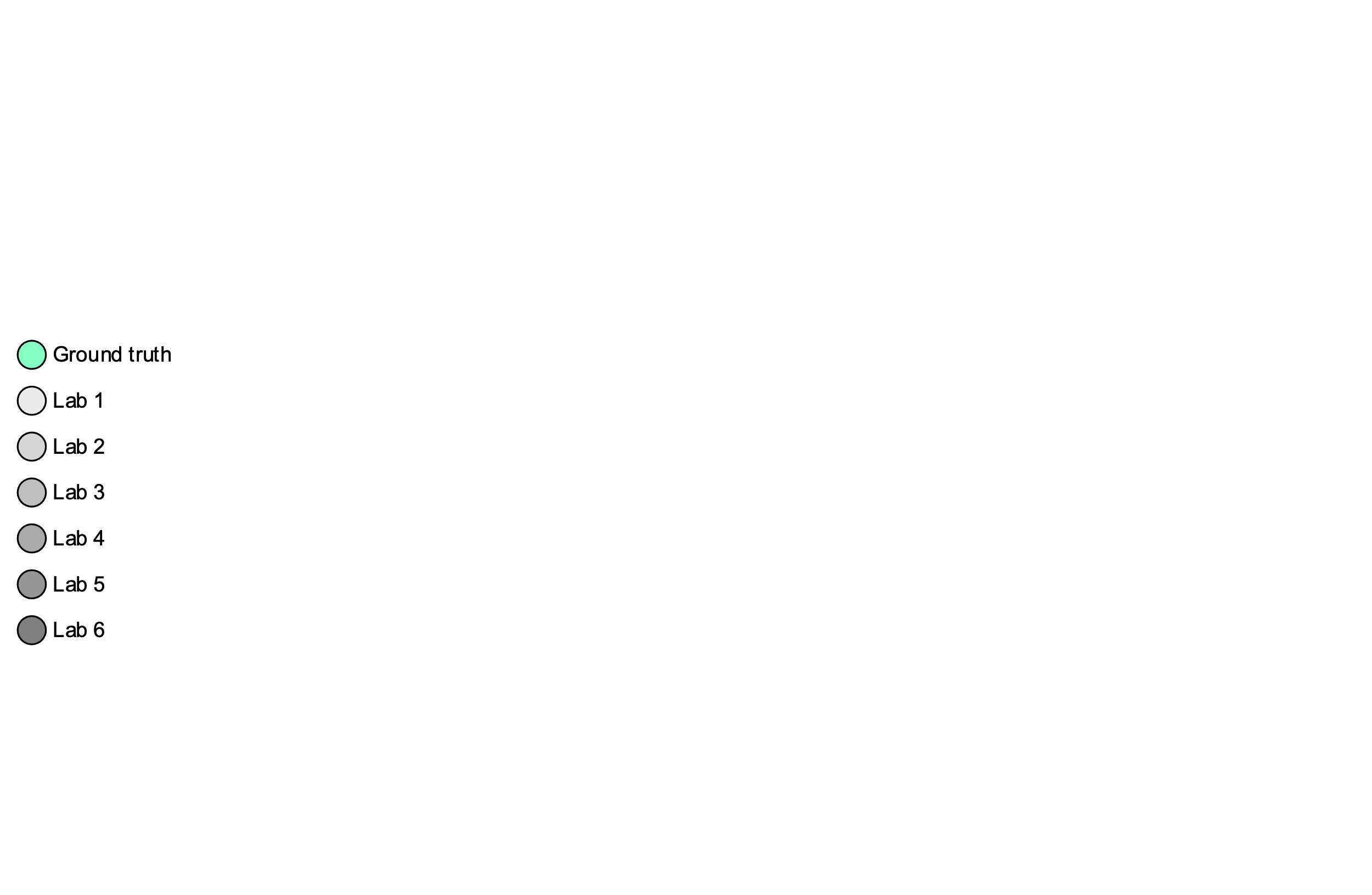

| **(b)** | **Avg. No. of Samples with Distance to GT** | **Max. cgMLST Allele Distance** | **Avg. cgMLST Allele Distance (SD)** | **Max. Missing cgMLST Alleles** | **Avg. Missing cgMLST Alleles (SD)** | **Avg. Ns Called** |
| --- | --- | --- | --- | --- | --- | --- |
| GT |  |  |  | 9 | 9.75 (1.63) |  |
| SUP m4.3 | 2 | 45 | 14.42 (16.75) | 73 | 25.21 (21.16) | n.a.* |
| SUP m4.3 + Polisher | 2 | 9 | 2.46 (2.90) | 158 | 58.63 (59.65) | 65.08 |
| SUP m5.0 | 1.83 | 3 | 0.71 (0,90) | 20 | 9.25 (3.35) | n.a. |
| SUP m5.0 + Polisher | 0.67 | 2 | 0.29 (0.44) | 39 | 14.08 (8.36) | 9.38 |
| * n.a. – not applicable | | | | | | |

**Supplemental Figure 3** (a) Minimum spanning tree of *M. morganii* (SUP m4.3, Medaka 2.0) cgMLST data without ONT cgMLST Polisher. Distances are based on cgMLST scheme of *M. morganii* (2,462 genes), pairwise ignoring missing values. The values on the connecting lines indicate the number of allelic distances between the connected isolates. Hybracter hybrid assemblies used as ground truth (green). (b) Error analysis results averaged over all sequenced ring trial isolates (n=96) with Dorado basecalling models SUP 4.3 and 5.0 with and without ONT cgMLST Polisher analysed.

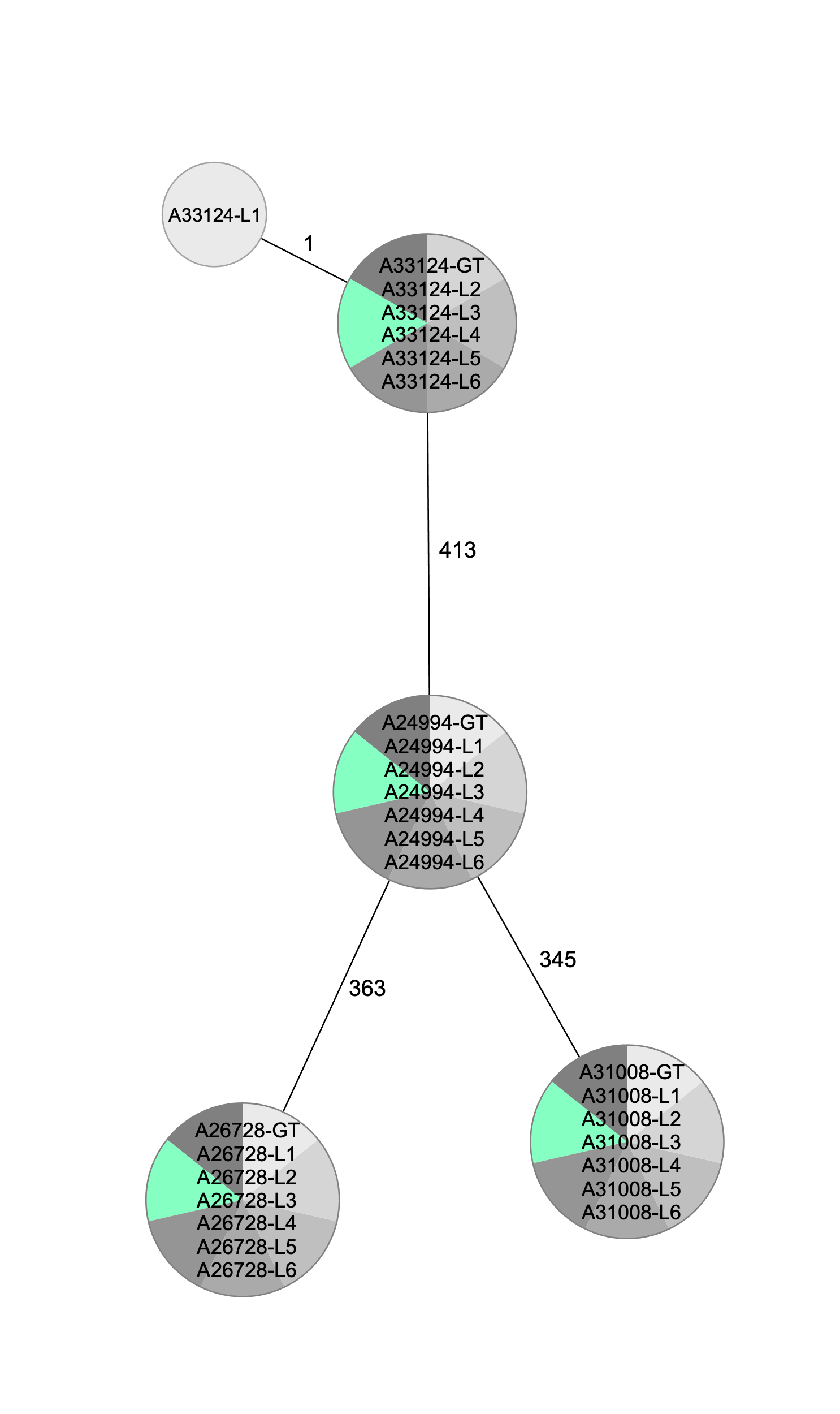
**(a)**

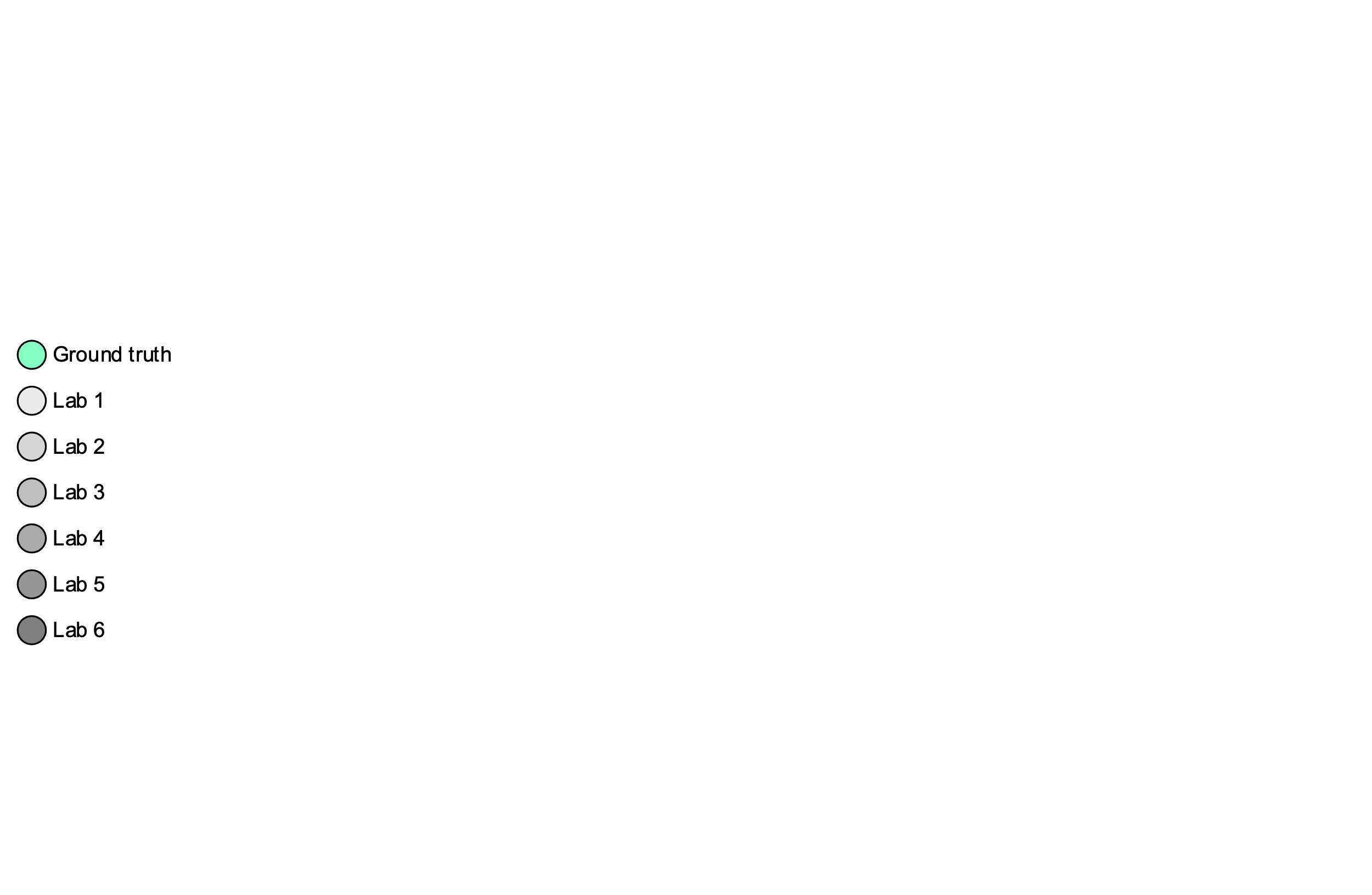

| **(b)** | **Avg. No. of Samples with Distance to GT** | **Max. cgMLST Allele Distance** | **Avg. cgMLST Allele Distance (SD)** | **Max. Missing cgMLST Alleles** | **Avg. Missing cgMLST Alleles (SD)** | **Avg. Ns Called** |
| --- | --- | --- | --- | --- | --- | --- |
| GT |  |  |  | 21 | 14.75 (8.10) |  |
| SUP m4.3 | 0.17 | 1 | 0.04 (0.08) | 21 | 14.96 (8.22) | n.a.* |
| SUP m4.3 + Polisher | 0 | 0 | 0.00 (0.00) | 22 | 15.17 (8.42) | 1.04 |
| SUP m5.0 | 0 | 0 | 0.00 (0.00) | 21 | 14.79 (8.13) | n.a. |
| SUP m5.0 + Polisher | 0 | 0 | 0.00 (0.00) | 22 | 14.96 (8.31) | 1.92 |
| * n.a. – not applicable | | | | | | |

**Supplemental Figure 4** (a) Minimum spanning tree of *E. faecium* (SUP m4.3, Medaka 2.0) cgMLST data without ONT cgMLST Polisher. Distances are based on cgMLST scheme of *E. faecium* (1,423 genes), pairwise ignoring missing values. The values on the connecting lines indicate the number of allelic distances between the connected isolates. Hybracter hybrid assemblies used as ground truth (green). (b) Error analysis results averaged over all sequenced ring trial isolates (n=96) with Dorado basecalling models SUP 4.3 and 5.0 with and without ONT cgMLST Polisher analysed.
